# Effects Of Video Games On Executive Control, Aggression and Gaming Motivation

**DOI:** 10.1101/2021.08.15.456380

**Authors:** Akshay Dixit, Divya Sinha, Hemalatha Ramachandran

## Abstract

With the advancements of computer technology and accessible internet, playing video games has become immensely popular across all age groups. Increasing research talks about the cognitive benefits of Video Games. At the same time, video games are stereotyped as an activity for the lazy and unproductive. Within this backdrop, our study aims to understand the effect of video games on Executive control (Visual Scanning and Visual Perception), Aggression, and Gaming Motivation.

Twenty non-gamers were selected and divided into two groups: Action Video Game Players (AVGP) and Non-Action Video Game Players NAVGP). We used two computerized tests: Gabor Orientation Identification Test and Visual Scanning Test (to assess visual perception and visual scanning, respectively) and two questionnaires (to assess aggression and gaming motivation). We found an improvement in visual perception as well as visual scanning following video game training in AVGPs. Interestingly, aggression did not increase with an increase in video game exposure. We also found insignificant changes in gaming motivation after the training, except for self-gratification motives.

Cognitive improvements do not relate to action video games alone, but non-action video games also show promising results to enhance cognition. With better timed and controlled training with video games, aggression as a prospective consequence of video game exposure can also be controlled. We propose targeted video game training as an approach to enhance cognition in non-gamers.

## INTRODUCTION

Video games have become an essential part of our life. People from almost every age group indulge in video games spaced over several genres. There are more than 2.5 billion active video gamers from all over the world (Wijman, 2019). Today’s gamer shows the traits of fortitude and efficiency and is also dexterous and innovative. Video games are designed for players to actively engage with cognitive systems and for these systems to, in turn, react to players’ cognitive demands (Granic, et al., 2014).

Action Video Games are one of the most played genres of Video Games. Studies have suggested that Action Video Games are involved in the development of numerous cognitive abilities. Significant evidence comes from training studies, especially in FPS type of video games, which are supposedly violent and action-packed, and not from simple puzzle games (Green and Bavelier, 2012). Playing Action Video games has been linked with a variety of enhancements like a reduction in response time of visual search rate (Green and Bavelier, 2003), a reduction of the attentional blink (Green and Bavelier, 2006), better change detection, and betterment in the number of items that can be tracked simultaneously (Green and Bavelier, 2006).

Research has also demonstrated that the skills learnt while playing action video games are retained even after the participant has lost exposure to the video game for up to one year (Bejjanki, et al., 2014).

Visual Scanning refers to the ocular strategies employed to explore various classes of visual stimuli like faces, objects or scenery. It is the ability to efficiently, quickly and actively look for information relevant to your environment.

Visual Perception is the processing of visual stimuli that makes sense of the physical world and its relationships to the body. It is a dynamic process that integrates all of the other senses (Bütün, et al. 2015).

Visual features of objects such as luminance, colour, orientation, motion, direction, and velocity are all registered pre-attentively (Wolfe, 1998). This pre-attentive processing may also decipher complex three-dimensional configurations (Enns and Rensink, 1990). Video games are inherently immersive. The scenes and the layout are incredibly dynamic and real life based. It becomes essential to read these visual stimuli by employing perceptual mechanisms of the brain.

Motivation is defined as the force that drives people to take action (Schiffman, et al., 2010). Different types of motivation may lead to various cognitive, behavioral and affective outcomes (Vallerand, 2001). Video games pull people of almost all ages into virtual world environments, making them work effectively into achieving meaningful goals, persevere in the face of multiple failures, celebrating the rare moments of triumph after the successful completion of tasks (Ferguson and Olson, 2013).

Human aggression can be thought of as any behavior that is directed to another individual and is carried out with an immediate intent to cause harm. Many video games contain a high amount of graphic and justified violence, especially shooter video games (Haninger and Thompson, 2004). A research study showed that playing violent video games affects the gamer’s real life, e.g., the confrontation with aggression in video games led to aggressive behavior in the real world (Gentile and Gentile, 2007).

Consecutively, a vast majority of research is aimed at looking at the negative and downsides of video games like depression, violence and gaming addiction. The value of such studies is important; however, a more balanced perspective is needed. Considering the potential benefits of playing video games is vital because the nature of these games has changed dramatically within the recent decade. It has become more sophisticated, diverse, realistic and social in nature. While Covid-19 has given us an opportunity to explore social media including time, consuming video games has become one of the ways to utilize this time. A study conducted by Bar and Copeland-Stewart (2021) highlighted an increase in gaming time in 71% respondents while 58% said that gaming has impacted their well-being. Interestingly, majority of respondents showed a positive impact.

Action video games possess in-game content and tasks that require the player to put Executive Control into practice to excel in the game. Therefore, a regular practice of playing the game should also train the players in these cognitive abilities. However, non-action video games do not contain such in-game tasks that require the player to use these abilities rigorously. Out of the many cognitive abilities pertaining to executive control, we assess visual perception and visual scanning. We hypothesized that the Action Video Game players would show an improvement in these abilities. Additionally, we also hypothesized that the Action Video Game players would show increasing aggression levels as the duration of video game exposure increases. Finally, we envisioned seeing an increase in gaming motivation post the training period.

We use a video game training study to approach our hypothesis; primarily because of its success with previous research. With a dynamically changing gaming environment, it becomes crucial to study the effects of video games on cognition to develop new and targeted video games, which may specifically help in the development of cognitive abilities and minimize the harm they may cause.

## METHODOLOGY

### Sample

23 Individuals participated in the longitudinal study. However, two participants dropped out of the study because they couldn’t complete the training period, and one dropped out due to personal health issues. One subject was dropped out of visual scanning experiment due to user induced confounds. Consecutively, the sample size was 20 consisting of males and females in equal numbers. All the participants had little or no exposure to video games. The participants filled in a demographics questionnaire and gave their consent for the study using a consent form. The participants recruited for the study were healthy adults in the age group of 18-25 years. All the participants had normal or corrected 6/6 vision. Only those people were recruited who had little or no exposure to video games.

Participants were encouraged to refrain from participating in any kind of gaming outside the training session. Participants post screening were divided into two groups: Action Video Game Players (AVGP) and Non-Action Video Game Players (NAVGP).

### Instruments

1. Games: We used two video games for training in our study.

a. Call of Duty: Black Ops (Treyarch, 2010) - Campaign mode with ‘recruit’ level of difficulty.
b. The Sims 2 (Maxis Redwood Shores Amaze Entertainment, 2004)

2. Computer-based tests: We used two tests as the pre-training and post-training tests to access executive control.

a. Gabor Orientation Identification Task (Dosher and Lu, 1999): The task followed a staircase design (Figure 1). It consisted of a Gabor patch which was rotated 10 degrees in a clockwise or anticlockwise direction. These Gabor patches were sandwiched between two levels of external noise, which were of Guassian type. Eight levels of external noise were used to sandwich the Gabor patch, as shown in Figure no. 2. The participants had to perceive the direction of rotation of the Gabor patch that was flashed in between the noise levels and respond by pressing either the ‘LEFT’ key for anticlockwise rotation or the ‘RIGHT’ key for clockwise rotation. A coloured dot appeared at the end of each trial to give the participant instant feedback—a green dot for correct response and a red dot for an incorrect response. The task consisted of a total of 128 trials, which was preceded by a trial sequence. We recorded accuracy as a measure of Visual Perception.
b. Visual Scanning Task: The distractors used in this task were letters. The participants had to focus on the fixation dot and actively scan the field of view to locate either of the two targets-the letter ‘F’ or ‘H’. The participants responded by pressing the ‘LEFT’ key for F and the ‘RIGHT’ key for H. A green dot was presented at the end of a trial indicating a correct response and a red dot was presented indicating an incorrect response. We used three paradigms in the task (Figure 3). The first paradigm consisted of 48 distractors and a target followed by the second paradigm consisting of 7 distracters and a target. The third paradigm again consisted of 48 distractors and a target. The total number of trials was 120. We recorded reaction time as a measure of Visual Scanning.

**Figure 1:**
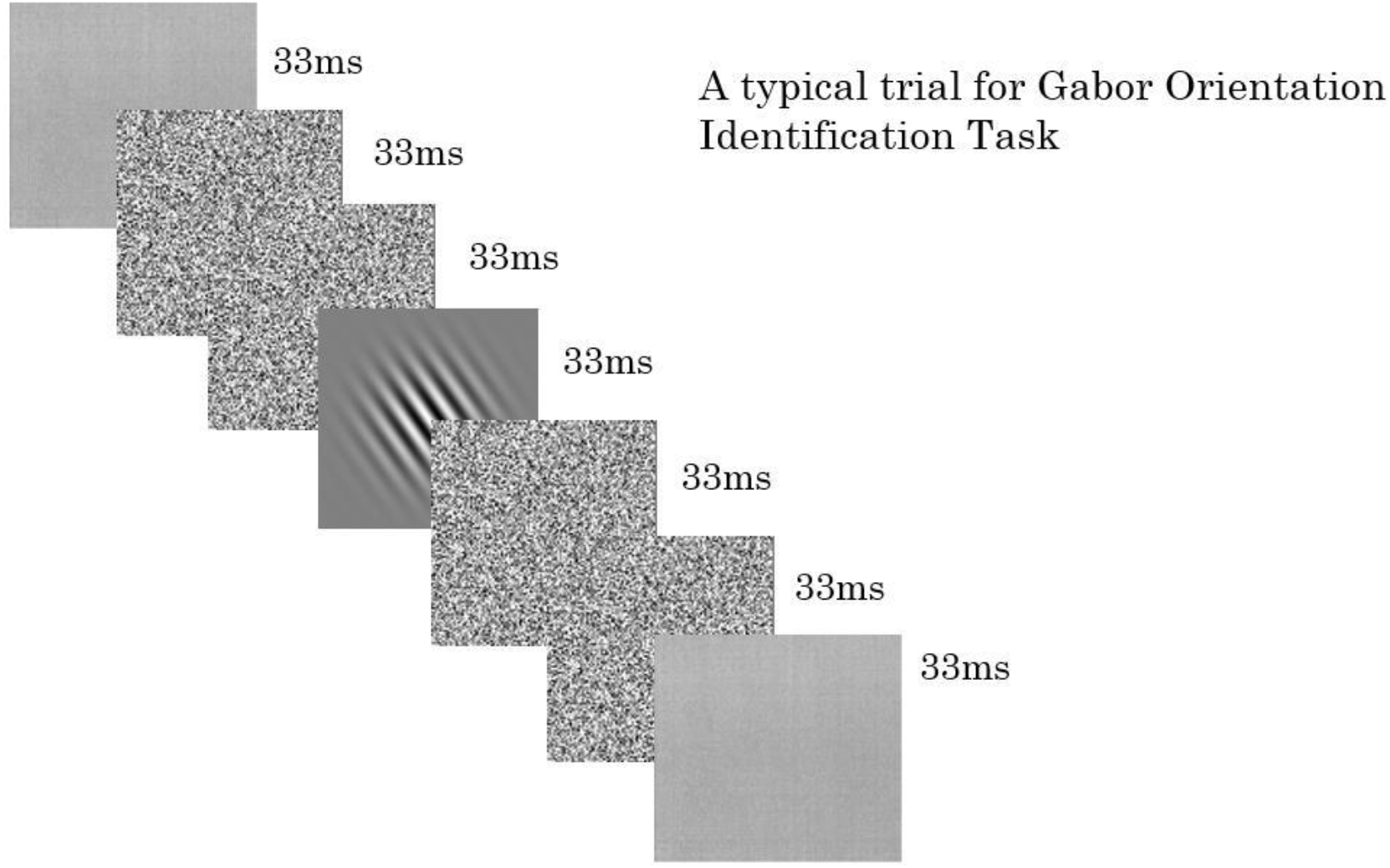
A typical trial for the Gabor Orientation Identification Task

**Figure 2:**
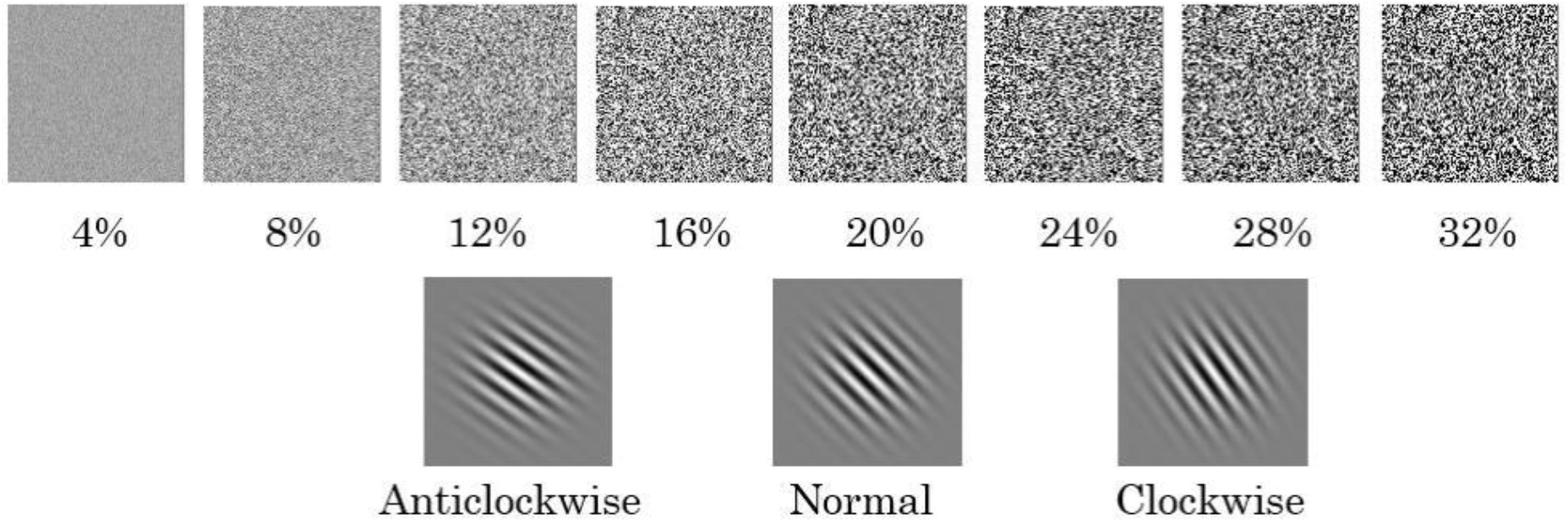
Depiction of the external noise levels and the Gabor Patches used in Gabor Orientation Identification Task

**Figure 3:**
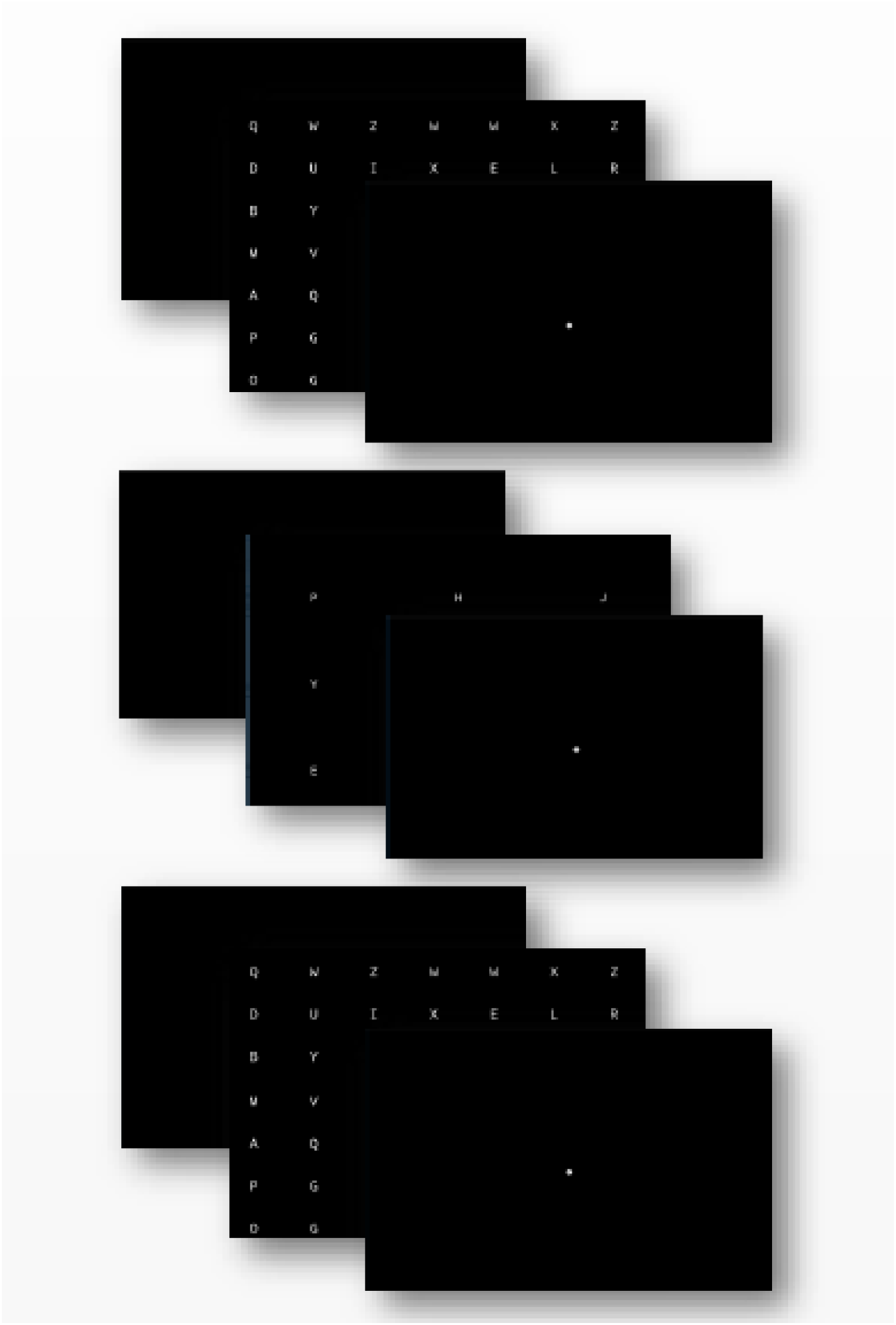
Flow of the Visual Scanning Task

### Questionnaires

We used two questionnaires, one to assess aggression and another to assess gaming motivation.

a. Buss-Perry Aggression Questionnaire (Buss and Perry, 1992): The participants were asked to fill in the questionnaire on an excel sheet before and after the video game training period. The questionnaire consists of a total of 29 items. The participants had to choose from a 7-point scale where 1 being ‘extremely uncharacteristic of me’ and 7 being ‘extremely characteristic of me’.
b. Electronic Gaming Motives Questionnaire (Myrseth and Notelaers, 2017): The participants were asked to fill in the EGMQ before and after the video game training period on an excel sheet. The questionnaire consists of a total of 14 items. The participants had to choose from a 4-point scale where 1 being ‘almost never’ to 4 being ‘almost always’.

### Technical Apparatus

We used four Intel Core i3 based Windows 10 laptops throughout the study. OPENSESAME v3.1.9 *Jazzy James (*Mathôt, et al. 2012) was used to administer the pre and post training tests to the participants. The tests were set on a resolution of 1024×768 px. We used MS Excel to administer the questionnaires. We used MS Excel and SPSS for data analysis.

### Procedure

Potential participants were recruited through personal interaction and advertisements on social media. After recruitment, each was given a demographics form and informed consent. The demographics form also consisted of screening questions. After the successful screening, we administered the pre-tests, which includes the two computer-based tests and the two questionnaires. Each participant was assigned a video game, i.e., either an Action Video Game or a Non-Action Video Game. The participants played the assigned game for one hour each day for a total of ten hours over 15 days. In other words, they played the game for ten days. After the final day of video game training, the participants were administered with the post-tests, which included the two computer-based tasks and the two questionnaires. After successful participation throughout the study, the participants were rewarded with a certificate of participation.

## RESULTS

### Effect of Video Games on Executive Control: Visual Perception

We expected that accuracy as a measure of visual perception would significantly increase in AVGP in the post-training task as compared to the pre-training task. A significant difference was obtained in accuracy levels between the pre and post training Gabor Orientation Identification Task in AVGPs but not in NAVGPs. (Table 1).

**Table No. 1.** Significance testing (Paired t-tests): Results from Gabor Orientation Identification Task measuring accuracy (%) between pre and post training in AVGP and NAVGP.

| Group | Pre/post | N | Mean | Std. dev | t | P |
| --- | --- | --- | --- | --- | --- | --- |
| AVGP | pre | 10 | 72.59343 | 18.67583 | -2.995 | 0.015* |
|  | post |  | 85.93196 | 10.23445 |  |  |
| NAVGP | pre | 10 | 66.36523 | 15.60957 | -2.107 | 0.064 |
|  | post |  | 74.57099 | 16.13383 |  |  |

Significance testing at $\alpha = 0.05$

### Effects of Video Games on Executive Control: Visual Scanning

We also expected that the reaction time as a measure of visual scanning would significantly decrease in AVGP in the post-training task as compared to the pre-training task and the same will not significantly decrease in NAVGP. Figure 5 (A) shows a decrease in the reaction time in post training test; and it was significant (Table 2). But we obtained insignificant results for NAVGP.

**Figure 4.**
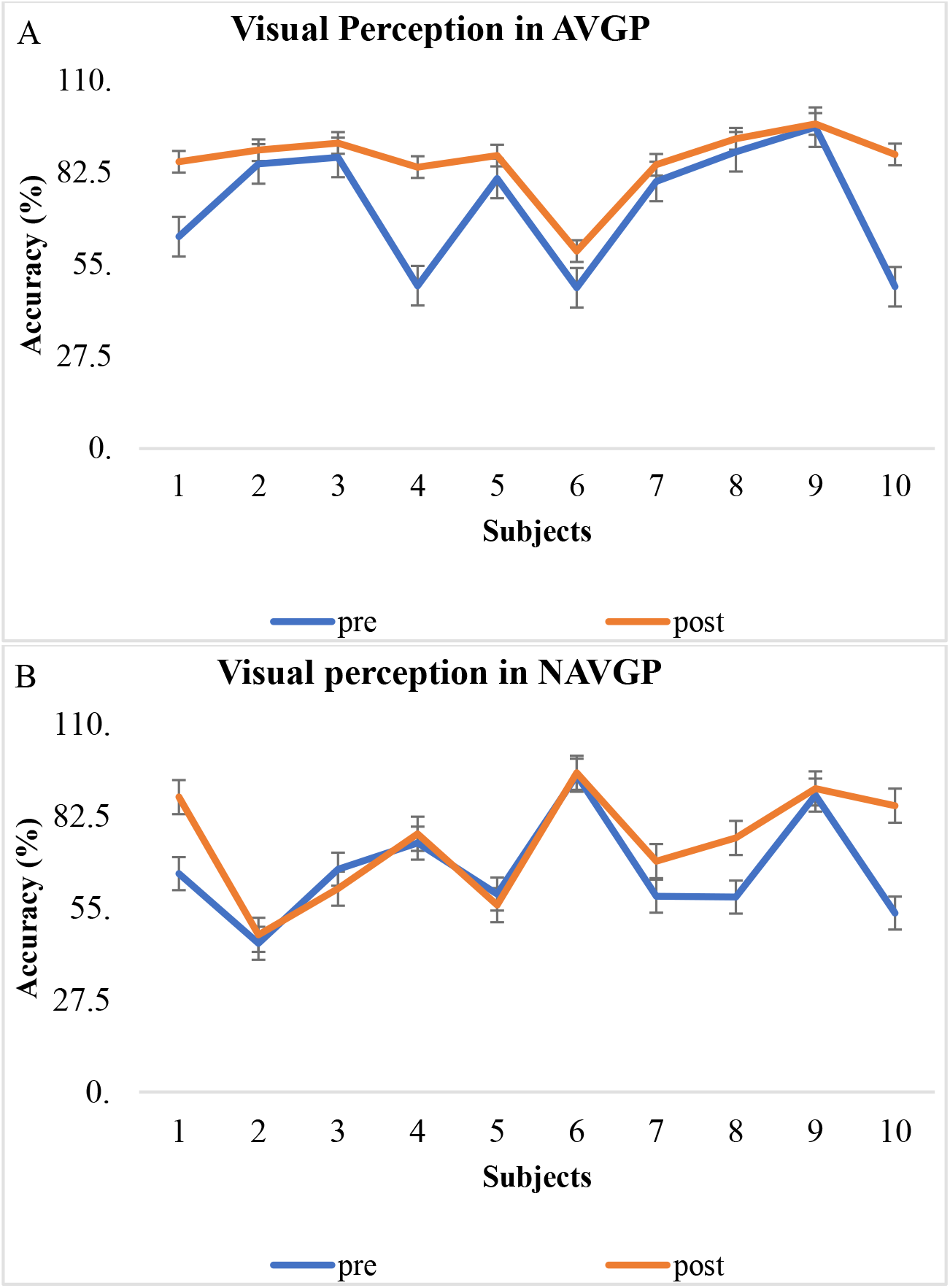
**(A)** Visual Perception in AVGP: We see that the accuracy levels were always higher in the post test as compared to pre-test. **(B)**Visual Perception in NAVGP: The post-test accuracy levels remain close to the pre-test accuracy levels with only a small increase or decrease throughout the training.

**Figure 5.**
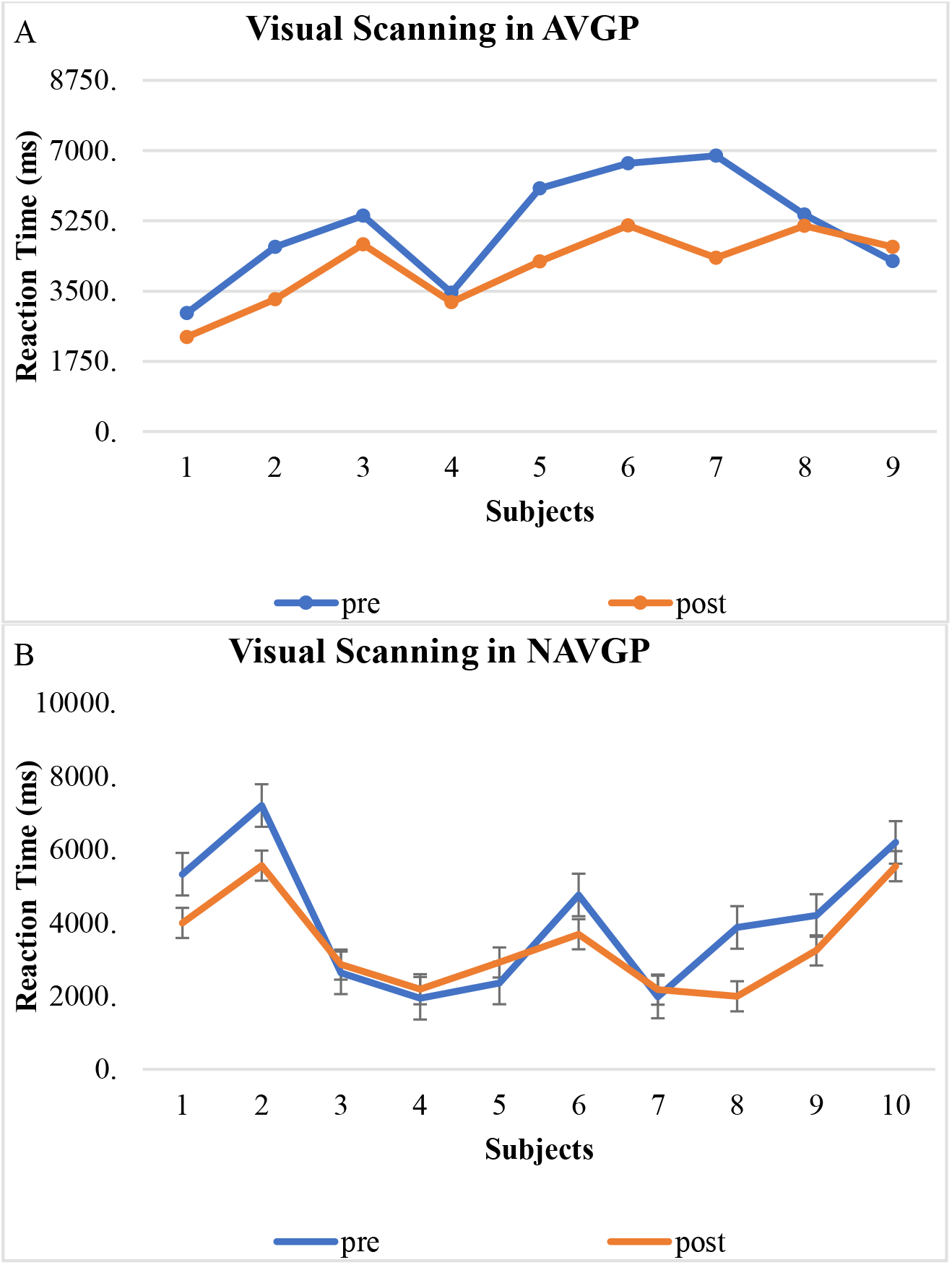
**(A)** Visual Scanning in AVGP: We see that the reaction time for post-test was generally less than the pre-test. **(B)** Visual Scanning in NAVGP: The reaction time for the post-test either increased or decreased as compared to the pre-test throughout the video game training.

**Table No. 2.** Significance testing (Paired t-tests): Results from Visual Scanning task measuring reaction time (ms) between pre and post training in AVGP and NAVGP.

| Group | Pre/post | N | Mean | Std. dev | t | P |
| --- | --- | --- | --- | --- | --- | --- |
| AVGP | pre | 9 | 5073.28 | 1369.424 | 3.194 | 0.013 |
|  | post |  | 4107.828 | 948.9715 |  |  |
| NAVGP | pre | 10 | 4045.316 | 1835.505 | 2.259 | 0.05 |
|  | post |  | 3416.479 | 1299.239 |  |  |

Significance testing at $\alpha = 0.05$

### Effects of Video Games on Aggression levels of AVGP and NAVGP

We were also interested in the effect of video games on the aggression levels across the study. We expected that there would be an increase in the aggression levels across both groups as the duration of exposure to video games increases. The average aggression score obtained from BPAQ shows a linear relationship by slightly falling as the duration of video game training increases (Figure 6). We found a negative correlation between the aggression levels and the duration of video game training; however, it was not significant (Table 3).

**Figure 6:**
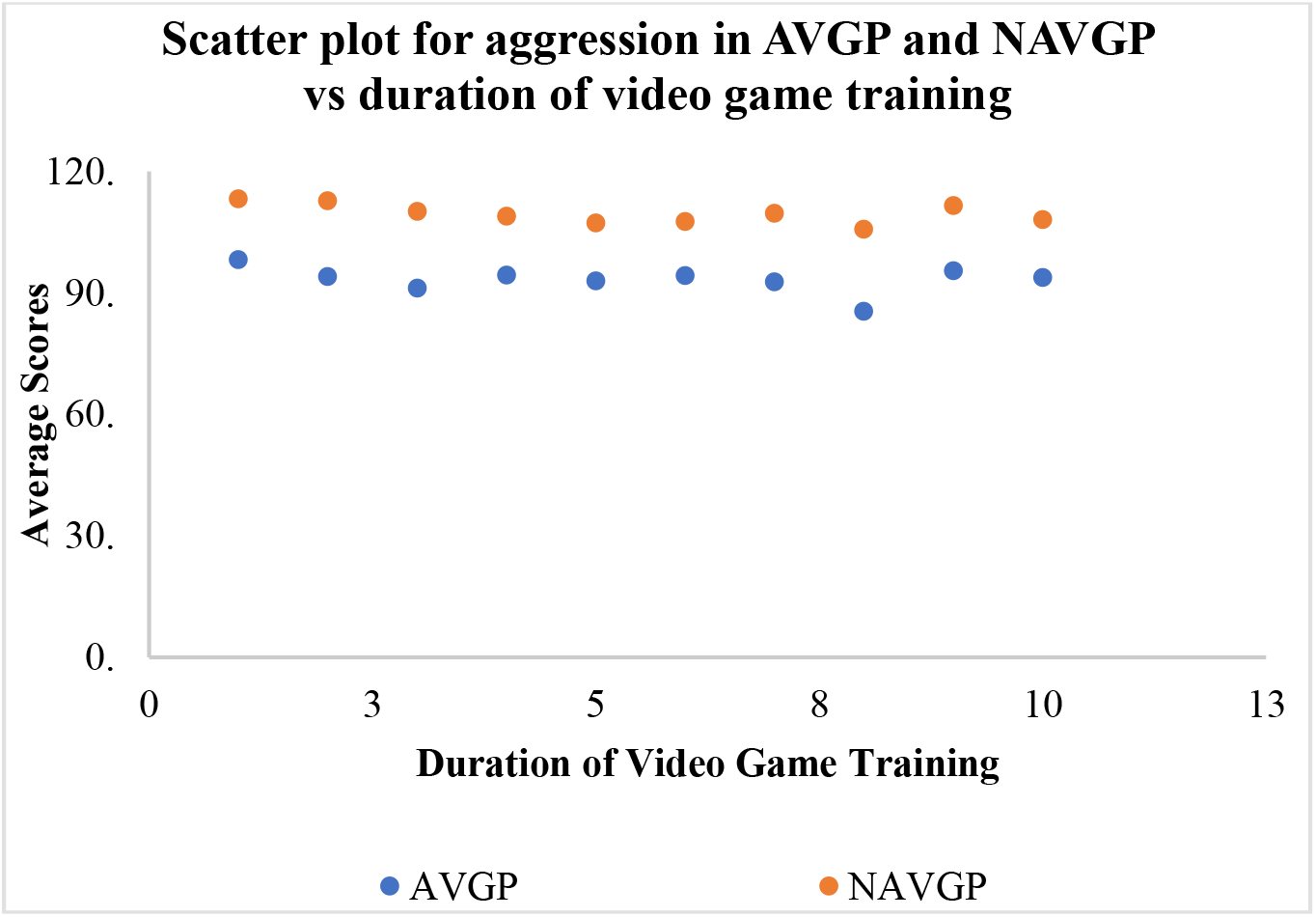
Scatter plot for aggression in AVGP and NAVGP vs. duration of video game training: We found a negative correlation between the aggression levels in both the groups vs. the duration of video game training. The figure shows that the correlation, though present, is not significant.

**Table No. 3:** Correlation analysis for aggression levels in AVGP and NAVGP across the training period.

|  |  | Days | NAVGP | AVGP |
| --- | --- | --- | --- | --- |
| Day | Pearson Correlation | 1 | -.557 | -.342 |
|  | Sig. (2-tailed) |  | .094 | .333 |
|  | N | 10 | 10 | 10 |
| NAVGP | Pearson Correlation | -.557 | 1 | .696* |
|  | Sig. (2-tailed) | .094 |  | .025 |
|  | N | 10 | 10 | 10 |
| AVGP | Pearson Correlation | -.342 | .696* | 1 |
|  | Sig. (2-tailed) | .333 | .025 |  |
|  | N | 10 | 10 | 10 |

\*. Correlation is significant at the 0.05 level (2-tailed).

### Effects of Video Games on Gaming Motivation of AVGP and NAVGP

We hypothesized that there would be a significant increase in all the four gaming motivation subscales in the post-test as compared to the pre-test in both the groups. To our surprise, only the self-gratification motives in AVGP increased significantly. The change in all the other subscales in both groups was insignificant (Table 4, Table 5).

**Table No. 4.** Significance testing (Paired t-tests): Results from EGMQ measuring 4 gaming motivation subscales of AVGP between day 1 and day 10.

|  | Day | Enhancement | Social | Coping | Self-gratification |
| --- | --- | --- | --- | --- | --- |
| Mean | day 1 | 3 | 1.766667 | 2.25 | 1.75 |
|  | day 10 | 2.866667 | 2.1 | 2.65 | 2.1 |
| Variance | day 1 | 1.432099 | 0.618519 | 0.652778 | 0.347222 |
|  | day 10 | 1.264198 | 0.569136 | 0.822222 | 0.502778 |
| P |  | 0.555445 | 0.138185 | 0.235256 | 0.024846 |

Significance testing at $\alpha = 0.05$ and $N=10$ .

**Table No. 5.**
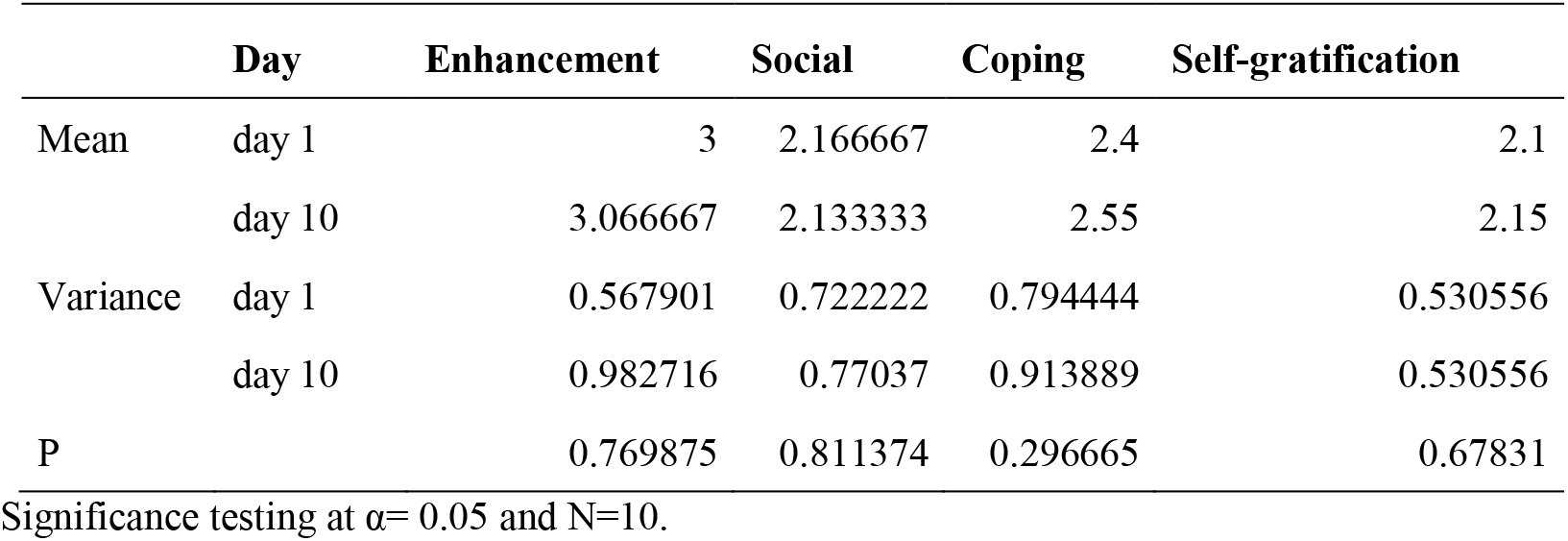
Significance testing (Paired t-tests): Results from EGMQ measuring 4 gaming motivation subscales of NAVGP between day 1 and day 10.

|  | Day | Enhancement | Social | Coping | Self-gratification |
| --- | --- | --- | --- | --- | --- |
| Mean | day 1 | 3 | 2.166667 | 2.4 | 2.1 |
|  | day 10 | 3.066667 | 2.133333 | 2.55 | 2.15 |
| Variance | day 1 | 0.567901 | 0.722222 | 0.794444 | 0.530556 |
|  | day 10 | 0.982716 | 0.77037 | 0.913889 | 0.530556 |
| P |  | 0.769875 | 0.811374 | 0.296665 | 0.67831 |

Significance testing at $\alpha = 0.05$ and $N=10$ .

## DISCUSSION

As all the participants had little or no exposure to video games, it was interesting to study the effects of video games on executive control: visual perception and visual scanning, aggression, and gaming motivation.

We found a significant increase in the accuracy of AVGPs after video game training. This suggests that action video games may have elements that train the player in better visual perception. However, we did not find such an improvement in NAVGPs, suggesting that the non-action video games may not be inclusive of the elements needed to enhance visual perception. Our results indicate the development of better perceptual templates involved in the enhancement of this cognitive ability (Bejjanki, et al. 2014).

We also found a significant decrease in reaction times of AVGPs. However, reaction time did not significantly decrease in NAVGPs. This result suggests that the action video games may enhance the visual scanning skills of the players but not non-action video games. Reaction Time and processing speed has been reported to enhance following action video game training (Oei and Patterson, 2013).

The duration and intensity of training has an important impact on effectiveness and feasibility of the training itself (Hempel et al., 2004). Researchers have used variable video game training periods. Cherney et al., (2014) used a one-hour training duration whereas Chandra et al., (2016) employed a fifty-hour training duration. Both reported significant improvement in cognition post training. Boot et al., used a twenty-hour training period. They found that there was no significant improvement seen in performance on visual tasks in participants who were trained on action video games for over twenty hours. This shows that the duration of the game intervention to see a significant improvement is still not understood.

Boot et al. (2008) recruited only non-gamer participants to the study. They had also suggested the comparison of the results of main group with an active control, i.e., a group that receives a game training intervention of a different type than that of the main group. In our study, the main group was the one playing an action video game and the non-action group was the active control.

The task designs employed in our study; however, slightly differ from those used by other studies reporting significant improvements (Dosher and Lu, 1999). Thus, the type of design could perhaps be playing a role in the failure of finding significant improvements. Other methodological errors—like multiple tasks influencing each other—might also lead to similar insignificant results across the groups.

The possibility of gender being a factor influencing the results is very low. Spence, et al, (2009) report that the learning trajectory of men and women is similar when acquiring basic spatial skills. They trained males and females of the same age group on an FPS video game and found similar results in both post training. Thus, the similar results found in the males, as well as females on the post training tasks in the current study, is in line with it, suggesting that both the groups may be equally trained on the visual tasks.

A negative correlation between the aggression levels and the duration of video game training suggests that as the duration of exposure to a video game increases, aggression levels go down. However, it is not significant as there is only a small deviation in the data points (Figure 6). This suggests that the stereotype that action video game play increases aggression in AVGP may not be fully valid. However, the fluctuations may also be as a result of external factors influencing the participant’s mood, and not the video game intervention.

Video games provide a dynamic environment—like switching levels or avatars—which demand players to flexibly adapt to new systems without experiencing frustration and anxiety (Granic et al., 2014). But these changes in the just learned game might displease some participants and possibly be a factor behind the small insignificant dip in enhancement motives in both the groups (Figure 7). A higher coping and self-gratification motives in action video game players (Figure 7, Table 4) suggest that the participants might indulge in gaming-related problems in the future (Myrseth, 2017). Identification of such motives for gaming is vital to understand what exactly differentiates between problem and non-problem gamers. And this information can be used to come up with effective interventions.

**Figure 7:**
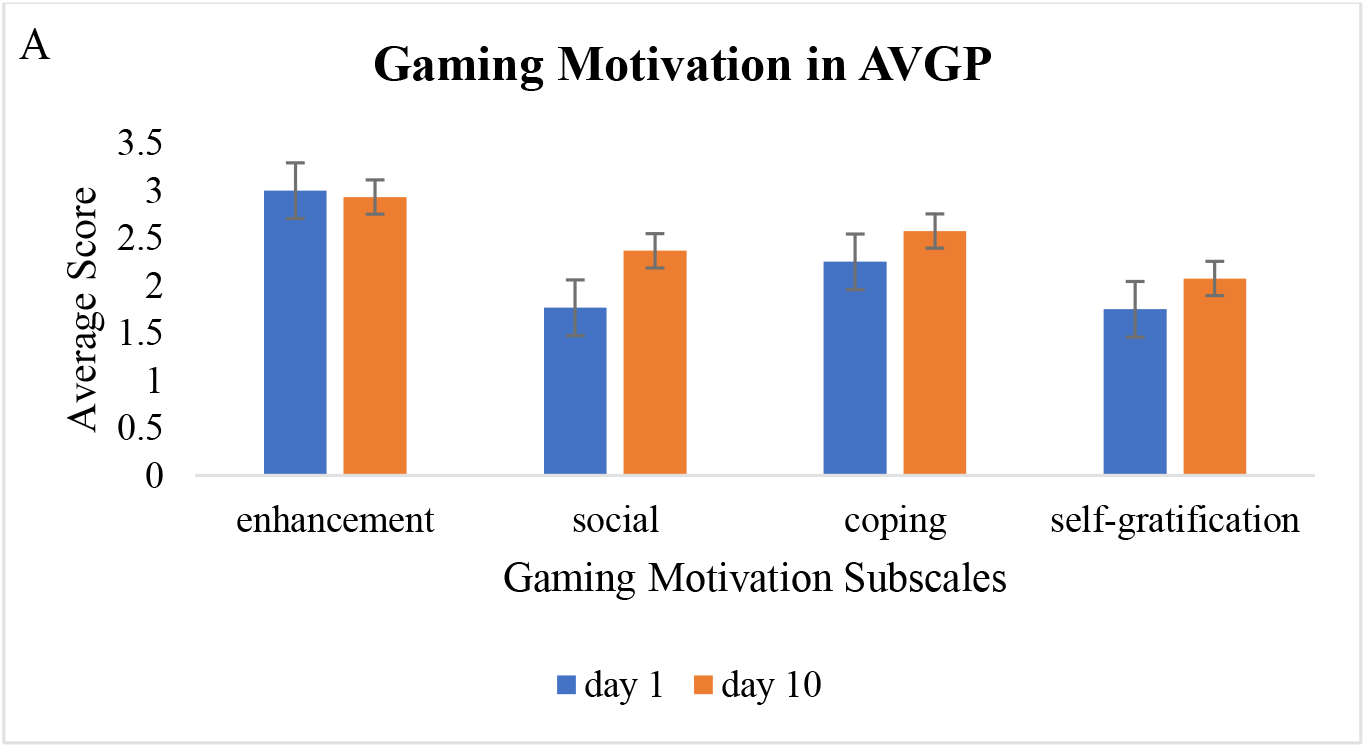
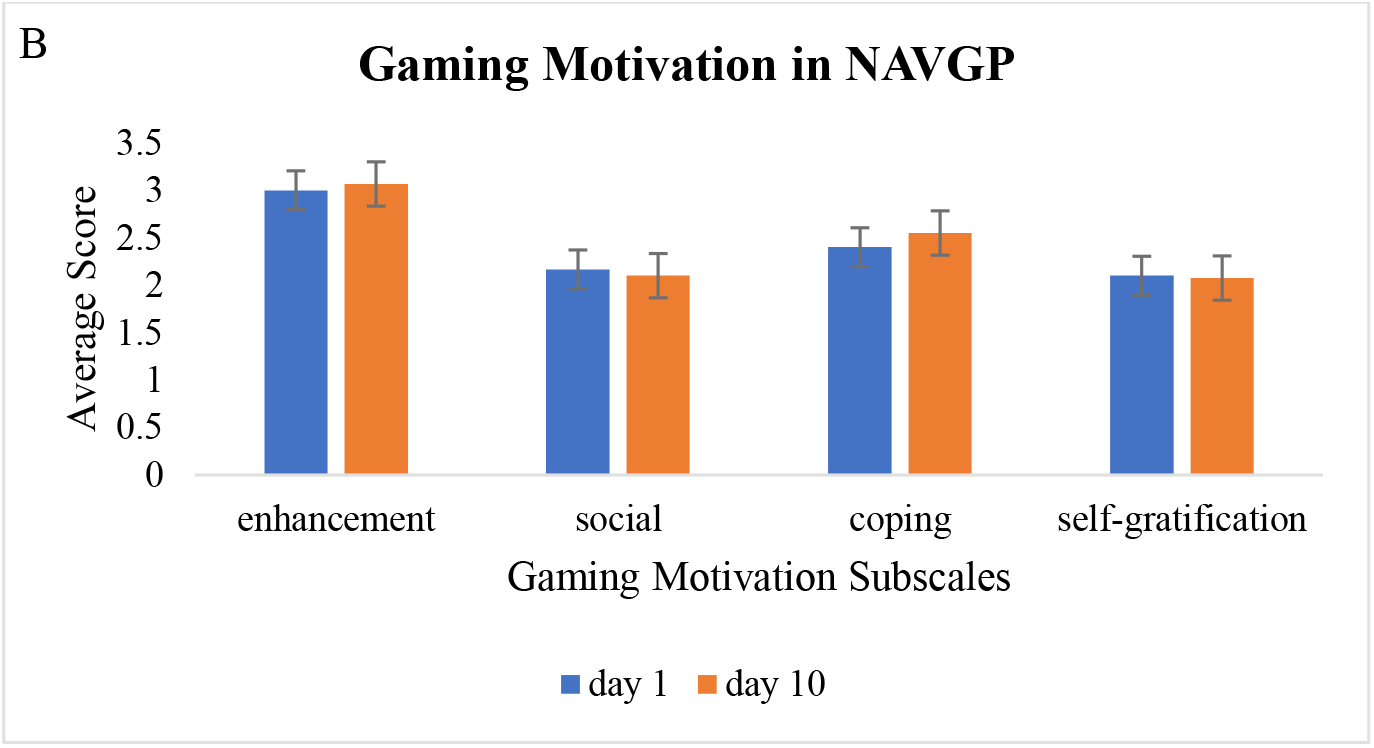
Gaming Motivation in AVGP and NAVGP. (A) We see an increase in the social, coping and self-gratification subscales of gaming motivation levels in the post-test as compared to the pre-test in AVGPs. We also observe a small fall in enhancement motives in the post-test (B). We see a small decrease in social and self-gratification motives in the post-test but a slight increase in enhancement and coping motives.

Overall, these results suggest that many video-games related cognitive improvement may not be due to training on broad cognitive systems like the executive control, but instead due to consistent utilization of specific cognitive process like visual scanning and visual perception during the game play.

### Limitations and Future prospects

The sample size per group was a major limiting factor in this longitudinal study. However, many studies have used a smaller sample size and managed to find significant results (Green and Bavelier, 2003; Bejjanki, et al., 2014). Also, the maintenance of a controlled environment could only be optimized and not guaranteed. This also can serve as another limiting factor during the training. Our study does not include robust statistical methods. Repeating the study using more powerful statistical methods will ensure better conclusive results.

It is essential to break down the video games into smaller parts and study the impacts to understand the underlying mechanisms of their engagement. To determine whether the results are consistent using our experimental design, a repetition of the study using more participants per group as well as targeted training duration is necessary.

Finally, it is important to caution that although video games show promising features of cognitive enhancement, they should not be treated as an ultimate tool to improve cognition. With increased virtual gaming during the pandemic, a healthy usage of video games is the need of the hour.

## Conclusion

With targeted exposure to video games, we can enhance executive control even in non-gamers. Participants reported a positive experience during the video game training regardless of their group. It can be said that regardless of the genre, video games lead to a positive experience. Thus, driving a gamer to exploit the sweet triumphs of video game rewards. We propose targeted video game training as a potential instrument that can be used in cognitive enhancement and neuropsychological rehabilitation. Hence, video games can be used as an effective medium to deliver content, which can potentially improve people’s cognitive abilities.

## Acknowledgments

We appreciate all those who participated in the study and helped to facilitate the research process. We also thank anonymous reviewers for their comments.

